# Cladoceran diversity build-up in newly created pondscapes

**DOI:** 10.64898/2026.02.20.706936

**Authors:** Bram Janssens, Robby Wijns, Maxime Fajgenblat, Pieter Lemmens, Thomas Neyens, Luc De Meester

**Author notes:** Corresponding author: Bram Janssens. Shared first authorship (Bram Janssens, Robby Wijns).

## Abstract

This study aimed to identify the most important drivers of cladoceran species richness build-up and species accumulation patterns in newly created ponds that collectively form entirely new pondscapes (networks of closely situated ponds in a landscape). A total of 26 newly created ponds across two newly established pondscapes were surveyed repeatedly (n=11) for key environmental pond variables and cladoceran community characteristics during the first three years after the pondscapes’ creation. The study ponds vary in surface area and maximum depth and cover a wide range of hydroperiods, from permanent to temporary systems with only a short hydroperiod. In total, 16 cladoceran species colonized the newly created ponds within the first three years of the pondscapes’ existence. Consistent with previous research, *Daphnia obtusa* was the first species to colonize ponds in both pondscapes. Macrophyte establishment was the most important local factor influencing the accumulation of cladoceran species over time, with vegetated ponds leading to faster species accumulation, likely because of the close association of chydorid species, *Simocephalus vetulus* and *Ceriodaphnia* spp with macrophytes. Our observations highlight the role of the establishment of macrophytes for cladoceran species richness build-up during early pond succession and offer valuable insights for designing resilient pondscapes that support rapid cladoceran species accumulation.

**Statements and Declarations:** The authors have no conflicts of interest to declare.

## Introduction

Over the past decades, biodiversity loss and climate change have reached an alarming rate (Sayer et al., 2025). Nature-based Solutions (NbS) are increasingly promoted as crucial strategies to protect biodiversity, mitigate the effects of climate change and support ecosystem services and Nature’s contribution to people (NCP) (Díaz et al., 2018; Keesstra et al., 2018; Mendes et al., 2020). The European Biodiversity Strategy for 2030 emphasizes the implementation of NbS to achieve its conservation goals (El Harrak & Lemaitre, 2023). Ponds harbor strong potential as NbS because they are ubiquitous globally and provide habitat for a variety of freshwater species, including many rare and endemic species (Dudgeon et al., 2006; Strayer & Dudgeon, 2010). Moreover, they serve as vital stepping-stone habitats that facilitate the dispersal of a wide range of species (Juračka et al., 2019), provide a wide diversity in ecosystem services and NCP, including carbon storage and climate mitigation (Postel & Carpenter, 1997; Cuenca-Cambronero et al., 2023; Lynch et al., 2023), and are relatively easy to construct at a relatively low cost (Cuenca-Cambronero et al., 2023).

Pond ecosystems are vulnerable to anthropogenic stressors, and are, due to their small size, largely overlooked by policymakers compared to larger aquatic systems such as large lakes and rivers (Bagella et al., 2016; Hill et al., 2021; Cuenca-Cambronero et al., 2023). As a result, these habitats are rapidly deteriorating, leading to steep declines in population sizes of their inhabitants and even the loss of the habitats themselves in recent decades (Dudgeon et al., 2006; Sayer et al., 2025). Both the reduction in quality and the reduction in their number have important impacts on biodiversity and landscape connectivity (Horváth et al., 2019). This highlights the urgent need for improved conservation measures targeting small water bodies (Biggs et al., 2016). To reach the full potential of ponds and pondscapes (networks of closely situated ponds in a landscape; Biggs et al., 2024; Bartrons et al., 2024) as NbS, increased know-how is needed on the management, restoration and creation of ponds and pondscapes.

Newly created ponds are typically non-equilibrium systems that display temporal variation in local environmental pond conditions (Williams et al., 2008; Coccia et al., 2016). Moreover, as early successional processes can have long-lasting effects on pond biodiversity and ecosystem functioning through priority effects and other legacy effects (De Meester et al., 2002; Chase, 2003), it is crucial to gain a deeper understanding of the processes at play in the early years upon pond creation.

Cladocerans are pivotal organisms in pond ecosystems (Leppänen, 2018; Ogorelec et al., 2021; Ebert, 2022), and are often used as study organisms in the context of metacommunity research (Cottenie & De Meester, 2003; Allen et al., 2011; Declerck et al., 2011; Gianuca et al., 2018). Cladocerans have been shown to rapidly colonize newly created ponds and their presence may boost the establishment of other species groups (Louette & De Meester, 2005), for instance by altering local pond conditions through top-down control of phytoplankton, contributing to a clear water state (Luecke et al., 1990; Effler et al., 2015). They provide an important food source for planktivorous fish and various macroinvertebrates, making them a crucial link between primary producers and higher trophic levels (Sterner, 2009). Cladocerans are therefore attractive as study organisms to identify the important drivers during early pond succession (Louette & De Meester, 2005).

In this study, we capitalize on a unique opportunity provided by two newly created pondscapes to study the cladoceran species richness build-up in newly created ponds. These pondscapes were constructed as part of a large nature restoration project in Eastern Belgium, conducted by Foundation Elzéard [*Elzéard*.*org*]. The aim of this study is to assess to what degree species richness build-up is determined by pond identity, pondscape identity, macrophyte presence, hydroperiod and other environmental pond variables (conductivity, chlorophyll a concentration, phycocyanin concentration, oxygen saturation, maximum depth, surface area, total nitrogen (TN), and total phosphorus (TP)).

We expect the studied ponds to display strong temporal variation in environmental conditions and species composition during the first three years of their existence and that the typical conditions of newly created ponds will be important in shaping early species accumulation patterns. If species accumulation trajectories are determined by species’ abilities to deal with these specific local conditions or if they are driven by differences in dispersal capacity between species, then we expect similar patterns across ponds within a pondscape. At the pondscape level, differences between pondscapes are expected if pond characteristics differ among pondscapes, if regional abundances of species are important and differ among pondscapes, or if regional species pools differ. If variation in local conditions across ponds is important in determining species accumulation trajectories, we expect a signal of pond or pondscape identity, but driven by the differences in local conditions rather than by identity of the pond or pondscape itself. Finally, if chance events linked to colonization events dominate the pattern, we expect highly diverging species accumulation curves that are not linked to local conditions or pondscape identity (Louette & De Meester, 2005).

## Materials and Methods

### Study site

Field sampling was conducted in 26 newly created ponds located in two newly established pondscapes situated in the East of Belgium: Rullen (n=16, 50°43’27.4” N 5°49’03.9”E) and Boffereth (n=10, 50°44’45.7” N 5°56’53.1”E) (Fig. 1). Each pondscape consisted exclusively of newly created ponds, with no previously existing ponds located between them. Additional information on the distances of all studied ponds to the closest existing (i.e. older, not newly created) regional pond is given in supplementary information (Table S1.1).

**Fig. 1:**
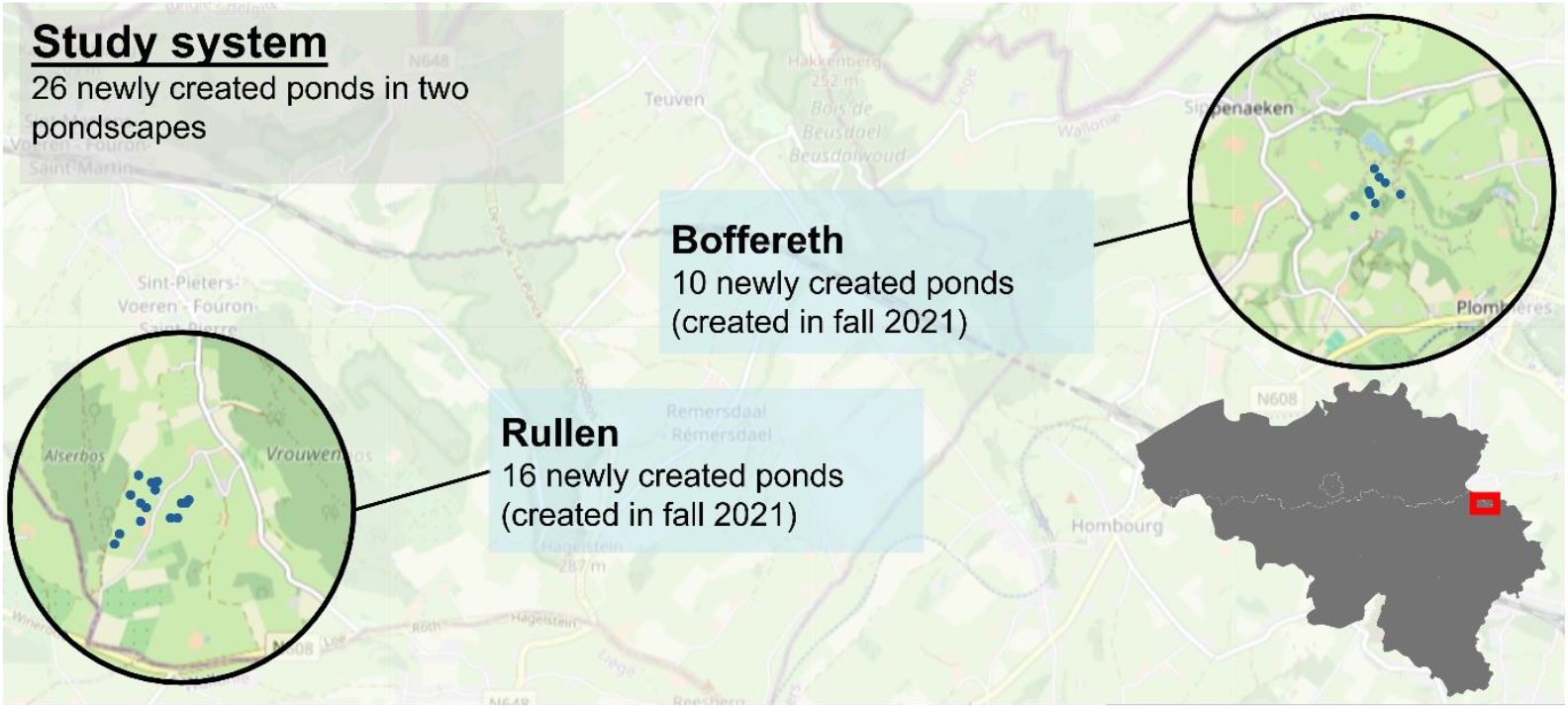
Geographical location of the study system consisting of newly created ponds in two newly established pondscapes. The inset map (bottom right) shows where the study system is located in Belgium (red rectangle).

Most ponds were created in November 2021, with the exception of one pond created in June 2022 (RUL16), and two ponds created in June 2023 (RUL1 & RUL2). During construction, the ponds were lined with a Dernoton® clay mixture and clay sourced from a local mine, precluding inoculation via resting eggs and allowing natural colonization. The ponds vary in surface area (3-27 m^2^), depth (up to 1.8 m), and hydroperiod, ranging from permanent ponds to ephemeral ponds that hold water only during the wettest seasons. More detailed information of these pond characteristics are given in Table S1. All ponds filled naturally through rainfall or groundwater seepage, with the exception of RUL3, 4, 5, and 6. These four ponds were manually filled using water from a local well, which we later found out contained *Daphnia obtusa* (Kurz, 1874), *Simocephalus vetulus* (O.F. Müller, 1776), and *Chydorus sphaericus* (O.F. Müller, 1776). Given that these species may have been introduced into these ponds, data of these ponds were not included in the analyses presented here; a parallel analysis of the dataset including these four ponds is given in supplementary information (SI2).

### Data collection

All ponds were monitored seasonally (four sampling campaigns a year: spring, summer, autumn and winter) to provide a temporally resolved picture of cladoceran diversity build-up over the first three years of the pondscapes’ existence. The first sample event occurred three to four months after pond creation. Given that we sampled all ponds during a three year period (November 2021– November 2024) and that some ponds were constructed in the years following the initial creation of the pondscapes (notably: one pond in Rullen created in June 2022 and two ponds in Rullen created in June 2023), these ponds were monitored over a shorter period than three years. At each of the 11 sampling events, the cladoceran community was sampled by collecting a depth-integrated water sample using a tube sampler (length 1.2 m; diameter 75 mm) at several sampling locations situated on a wedge-shaped grid containing eight cells. Water from each location within each pond was pooled and 30 L from the combined water was subsequently filtered through a 64 µm mesh filter. The zooplankton sample was preserved in glucose-saturated formaldehyde (4%) until further processing in the laboratory. During sampling, we paid special attention to prevent cross-contamination. Ponds were only entered when necessary, and most of them could be sampled directly from the bank due to the small size of the ponds. If entering the pond was nonetheless needed to get representative samples, waders were thoroughly rinsed (including the sole of the boots to prevent egg bank transfers between ponds) before entering a new pond. Between the sampling of different ponds, all equipment was thoroughly rinsed and disinfected with H2O2 (diluted 1:7) to prevent cross-contamination between the newly created ponds. For each pond sampled on a given day, we used a different cylindrically shaped mesh filter to concentrate the zooplankton.

Cladoceran specimens were identified to species level using a binocular stereoscope (Olympus SZX16) and identification keys of Amoros (1984) and Flössner (2000). We followed the same standardized counting protocol as De Bie et al. (2012). Before identifying cladocerans, formaldehyde was replaced with water. The entire sample was then examined using subsamples (1–10 mL, depending on the organic material present in the sample), which were transferred into a counting tray using a broad-mouthed micropipette, preventing cladocerans from floating at the surface. To maximize detection and to avoid underestimating species richness, the entire sample was thoroughly screened. This procedure ensured that all individuals present in the sample were considered, allowing us to compile an accurate estimation of cladoceran species richness data (presence/absence) for each pond and timepoint.

During each sampling occasion, a set of key local environmental pond variables was quantified using a depth integrated water sample from different parts of the pond. Chlorophyll a (µg/L) and phycocyanin (RFU) concentrations were measured using a handheld fluorometer (Aquafluor, Turner Designs), taking the mean of three replicate measurements. Maximum pond depth was measured using a measuring stick. Water transparency was quantified using a Sneller’s tube. Conductivity (mS/cm), pH and oxygen (mg/L) were measured directly in the pond using a HACH HQ 40d multimeter. To determine concentrations of TN and TP, expressed in milligrams per liter (mg/L), water samples were collected and frozen at −20°C for later analysis in the laboratory. Nutrient concentrations were measured after alkaline persulfate digestion (Koroleff, 1970) using a QuAAtro continuous flow analyzer (Seal Analytical). The percentage of pond surface covered with emergent, submerged and floating vegetation was visually estimated during each sampling event.

### Statistical analysis

A principal component analysis (PCA) was performed on scaled (standardized) data of local environmental pond variables (conductivity, chlorophyll a concentration, phycocyanin concentration, dissolved oxygen concentration, maximum depth, surface area, TN, and TP) to visualize the association between environmental variables, pondscape and time. A similar PCA analysis was carried out for both pondscapes separately to visualize potential differences in environmental associations between the pondscapes. For the maximum depth and the surface area, the highest value measured over the entire sampling period was used The PCA was conducted using the prcomp-function from the R package ‘stats’.

We used linear mixed models to study how the rate of community buildup, during the first three years of a pond’s existence, is modulated by pond identity, pondscape identity, macrophyte presence, hydroperiod and abiotic factors. We modelled the cumulative species richness, defined as the total number of observed species up to each sampling moment, as primary outcome to overcome imperfect species detection and seasonal variation in the observed instantaneous species richness. We modelled the cumulative species richness as a linear function of pond age (expressed in years) with an identity link function to approximate the rate of community buildup. Since the cumulative species richness of a pond is known to be zero at age zero, we omitted the model intercept and any other main effects other than pond age. Instead, we included pondscape identity (binary; Boffereth (0) and Rullen (1)), hydroperiod (fraction of sampling moments during which the pond was filled with water throughout the sampling period), macrophyte coverage (binary; defined as having reached >25% emersed or submerged plant coverage at least once throughout the study period; the 25% cut-off was chosen because macrophyte coverage rarely fell below this level once 25% coverage was exceeded, except for short-term declines caused by large water influxes; a 25% macrophyte coverage thus indicates persistent macrophyte establishment), two proxies of environmental pond conditions (standardized PC1- and PC2-coordinates extracted from the abovementioned PCA analysis, averaged over the entire study period per pond), and distance to an established pond (z-score standardized distance to the closest regional pond that existed before the pondscapes were created) as interaction terms with pond age, to estimate how these variables change the rate of community buildup. To account for unexplained pond-specific variation in the rate of community buildup, we included a pond-specific random slope for the effect of pond age. We fitted this model using the R package ‘brms’ (Bürkner, 2017), which relies on the probabilistic programming language Stan and which performs Bayesian inference through a Hamiltonian Monte Carlo MCMC algorithm (Carpenter et al., 2017). We used weakly informative and mildly regularizing standard normal priors on all regression parameters, the random slope scale parameter and the residual error parameter. We used trace plots and the potential scale reduction factor to assess model convergence, and used posterior predictive checks to ascertain goodness of fit (Gelman et al., 2013).

To evaluate the degree to which communities in species poorer ponds form nested subsets of communities in species richer ponds, we conducted a NODF analysis (Nestedness metric based on Overlap and Decreasing Fill; Almeida-Neto et al., 2008) based on presence-absence data in the third year, using the nestednodf-function from the package ‘vegan’ (Oksanen et al., 2013). To assess whether the observed nestedness was greater than expected by chance, we compared the observed NODF to a null distribution based on 1000 randomized matrices. Random matrices were generated using the permatfull-function from the package ‘vegan’ (Oksanen et al., 2013), considering four null model algorithms: free row and column totals (most liberal approach), free row totals and fixed column totals, fixed row totals and free column totals, and fixed row and column totals (most conservative approach). p-values were calculated as the proportion of randomized NODF values that were equal to or greater than the observed NODF.

All visualizations were created using the ggplot-function from the R package ‘ggplot2’ (Wickham, 2011). All statistical analyses and visualizations were carried out in R (version 4.4.3) (R core Team, 2025). All code and data to reproduce the analysis can be accessed through Github (https://github.com/BramJanssens1/Cladoceran_richness_accumulation).

## Results

### Environmental properties of newly created ponds

The first and second axis of the PCA ordination based on standardized local environmental variables jointly explained 49.4% of the variation between ponds throughout the first three years (Fig. 2). The first axis (eigenvalue=2.66) was primarily associated with variation in chlorophyll a concentration, conductivity, pH, TN, TP, and daytime oxygen concentration. The second axis (eigenvalue=1.79) was predominantly associated with phycocyanin concentration, maximum depth, and surface area. Separate ordinations for both pondscapes showed strong similarities in overall patterns between the pondscapes (e.g., the positive association between chlorophyll a concentration, phycocyanin concentration, oxygen concentration and pH – all associated with photosynthetic activity), but with a positive association between TP and TN in the Rullen pondscape, whereas they are rather negatively associated in the Boffereth pondscape (Supplementary Information, Fig. S1.1). Ponds exhibited the highest variability in their environmental pond conditions the first year after their creation, after which they converged into more stable and similar conditions.

**Fig. 2:**
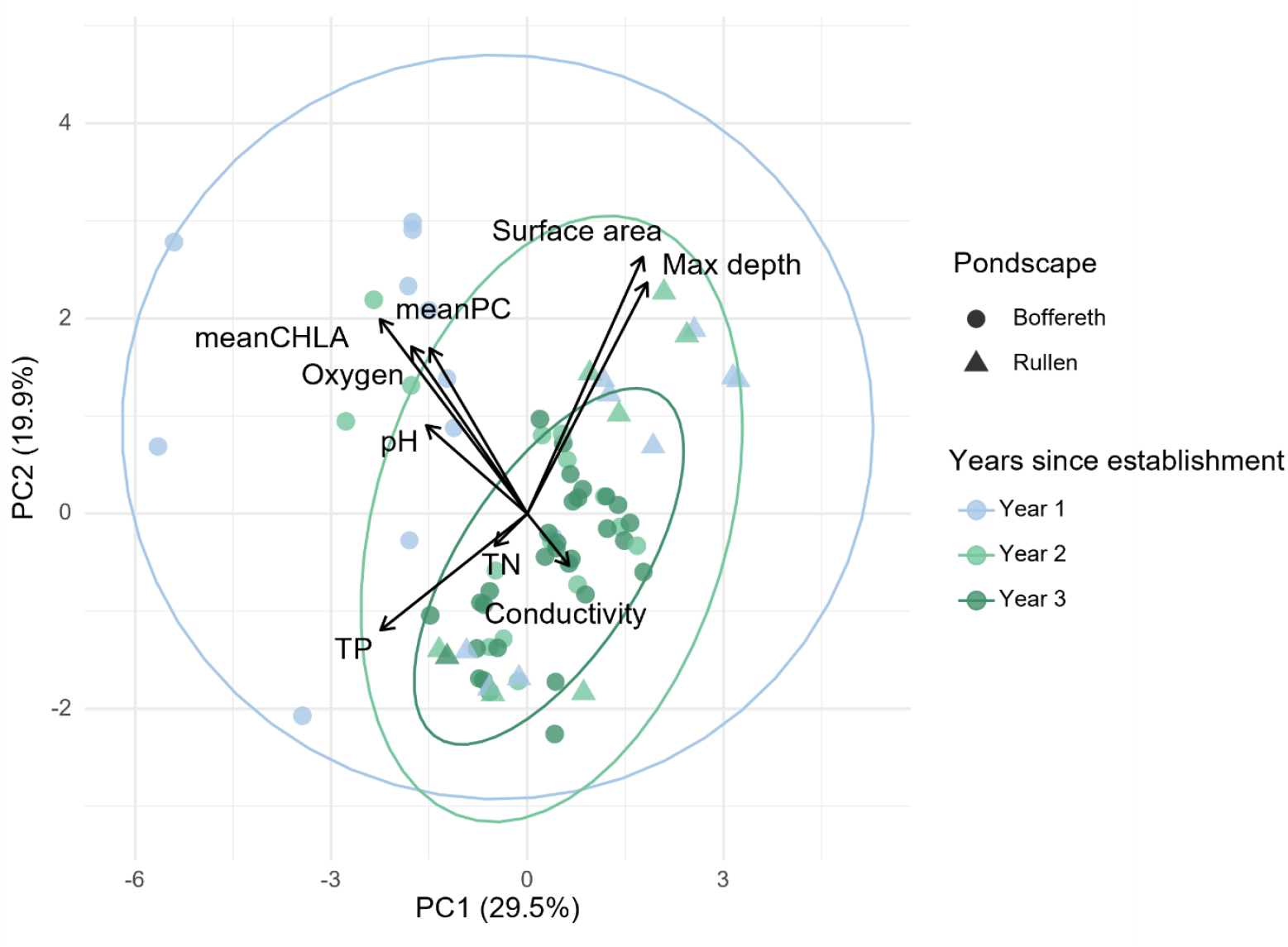
PCA biplot based on the environmental pond characteristics for each sampling moment, maximum depth and surface area during the first three years of their existence. Different years are highlighted using different 95% confidence ellipses. Pondscapes are visualized by different symbols (Boffereth: circles; Rullen: triangles).

### Cladoceran species richness build-up

The mean cumulative species richness clearly increased in all ponds in both pondscapes during the first three years after pond creation. Species accumulation rates were generally higher in Rullen compared to those in Boffereth (Fig. 3), and varied considerably between ponds within both pondscapes. When focusing on the overall accumulation patterns, we observed that, after an initial steep increase over time in the first months, species richness leveled off for nearly a year from month 7 onwards, to then increase again in the second half of the second year after pond creation (Fig. 4). BOF5 ultimately hosted the highest number of cladoceran species (n=12) by the end of the three-year period.

**Fig. 3:**
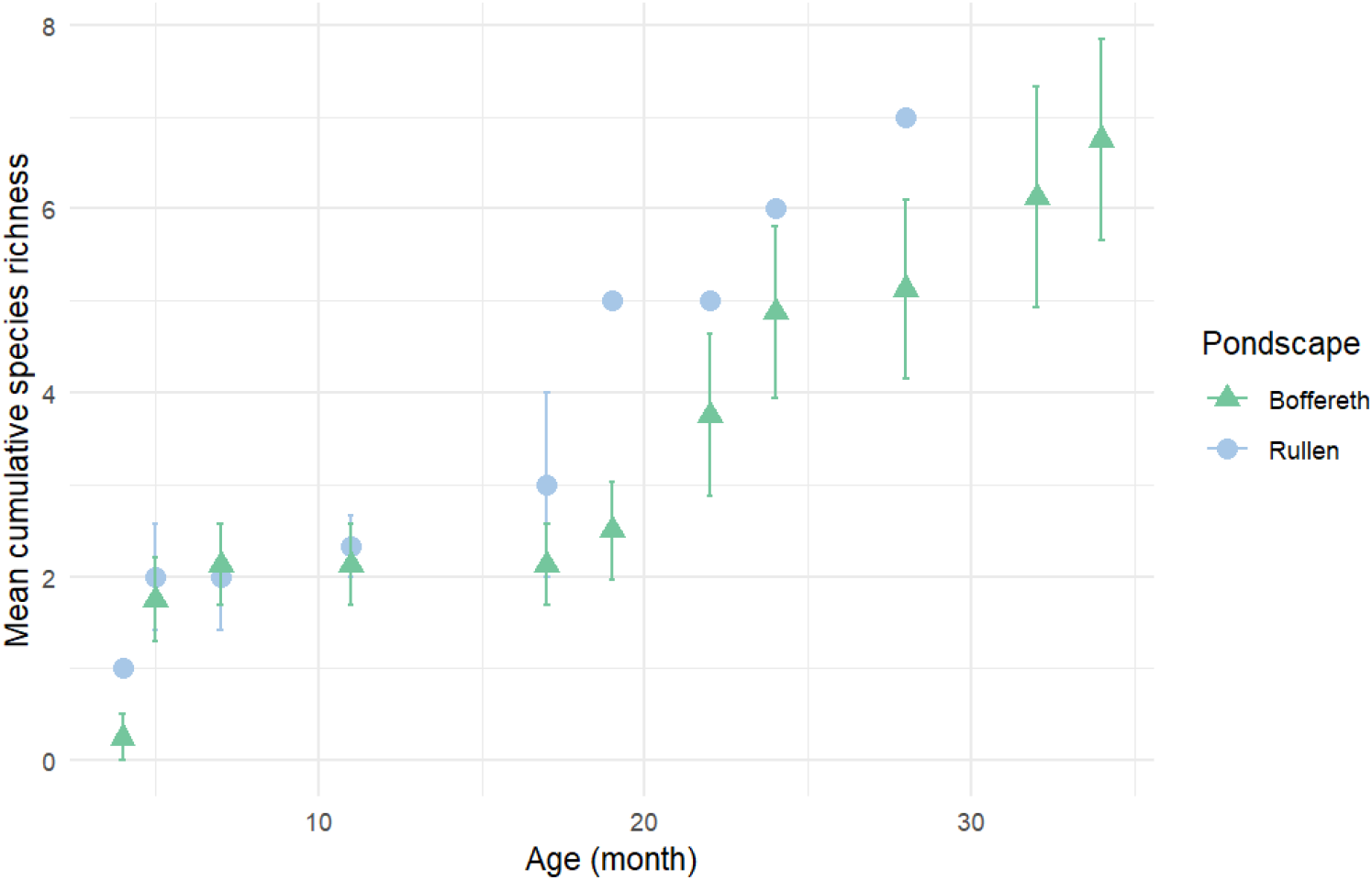
Average cumulative species richness of cladocerans during the first three years after pond creation for each newly created pondscapes: Boffereth (green triangles) and Rullen (blue circles). Values are averaged across ponds per pondscapes. Error bars indicate the 95% standard error.

**Fig. 4:**
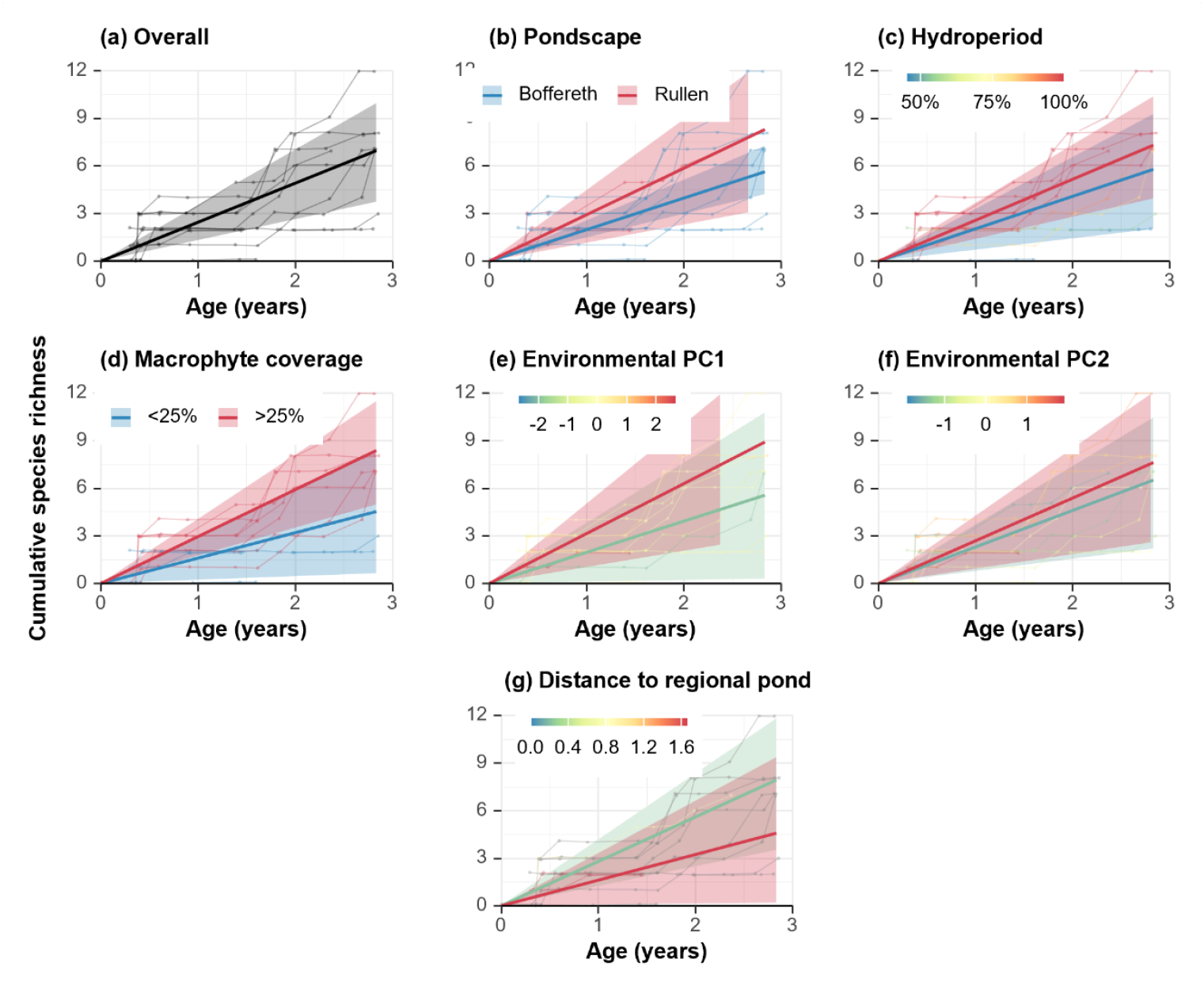
Estimated, conditional linear species accumulation trajectories as a function of pond age, all other conditions being held at the average value across the dataset. Individual points represent observed cumulative species richnes data across ponds and time points, with points belonging to the same pond being joined by thin lines. Estimated linear species accumulation trajectories are shown using thick lines, with shaded areas representing 95% credible intervals. (a) Estimated trajectory for the average pond. (b) Estimated trajectory for the average pond in the pondscapes Boffereth and Rullen. (c) Estimated trajectory for the average permanent and temporary pond. (d) Estimated trajectory for the average well- and less-vegetated pond. (e) Estimated trajectory for the average pond with a low and high local environmental condition as represented by the PC1 coordinate. (f) Estimated trajectory for the average pond with a low and high local environmental condition as represented by the PC2 coordinate. (g) Estimated trajectory for the average pond with a low and high distance to the closest regional pond.

Despite the observed difference in accumulation rate between ponds within pondscapes, we did not find strong statistical support for a direct effect of pondscape identity per se (16.4% posterior probability of an effect). An average pond (i.e. conditional on average characteristics, except for pondscape identity) in Rullen and Boffereth displays an estimated accumulation rate of 2.7 species per year (95% CrI [1.3; 4.0]) and 1.8 species per year (95% CrI [1.1; 2.6]), respectively (Fig. 4). We found strong statistical support that well-vegetated ponds feature higher accumulation rates than less-vegetated ponds (99.1% posterior probability of an effect). The average well-vegetated pond (highest recorded coverage of emersed or submerged macrophytes >25%) displays an estimated accumulation rate of 2.6 species per year (95% CrI [1.2; 3.9]), while the average less-vegetated pond displays an estimated accumulation rate of 1.3 species per year (95% CrI [-0.2; 2.7]) (Fig. 4). This corresponds to a difference in accumulation rate of 1.3 species per year (95% CrI [0.3; 2.3]). We also found moderately strong statistical support for a positive effect of hydroperiod (91.4% posterior probability). The average pond containing water during all sampling moments displays an estimated accumulation rate of 2.2 species per year (95% CrI [0.9; 3.6]), while the average pond containing water during half of the sampling moments displays an estimated accumulation rate of 1.7 species per year (95% CrI [0.5; 2.9]) (Fig. 4). This corresponds to an estimated difference in accumulation rate of 0.5 species per year (95% CrI [-0.3; 1.3]). We found the joint posterior distribution of macrophyte status and hydroperiod to be negatively associated, reflecting that macrophyte vegetations are better developed in ponds with longer than with shorter hydroperiods (Fig. 5). As a result of this collinearity, the model cannot fully disentangle the direct effect of macrophyte status and hydroperiod at a 95% certainty threshold. While we found strong and moderately strong statistical support for the marginal effect of macrophytes and hydroperiod, respectively, the available data does not allow to fully isolate the effect of macrophyte status, leading to a more uncertain estimate and moderately strong statistical support. We found 88.6% posterior probability that both variables are directly related to the species accumulation rate, but only a 10.6% posterior probability for a pure effect of macrophyte (i.e., excluding the shared effect with hydroperiod), and a 0.8% posterior probability for a pure effect of hydroperiod (i.e., excluding the shared effect with macrophyte status; Fig. 5). We did not find any evidence for the influence of local environmental conditions as summarized by the temporally averaged first and second principal component coordinates (80.9% and 68.5% posterior probability of an effect, respectively) (Fig. 4). We did not find strong statistical support for the effect of distance to the closest existing, regional pond (82.8% posterior probability of an effect).

**Fig. 5:**
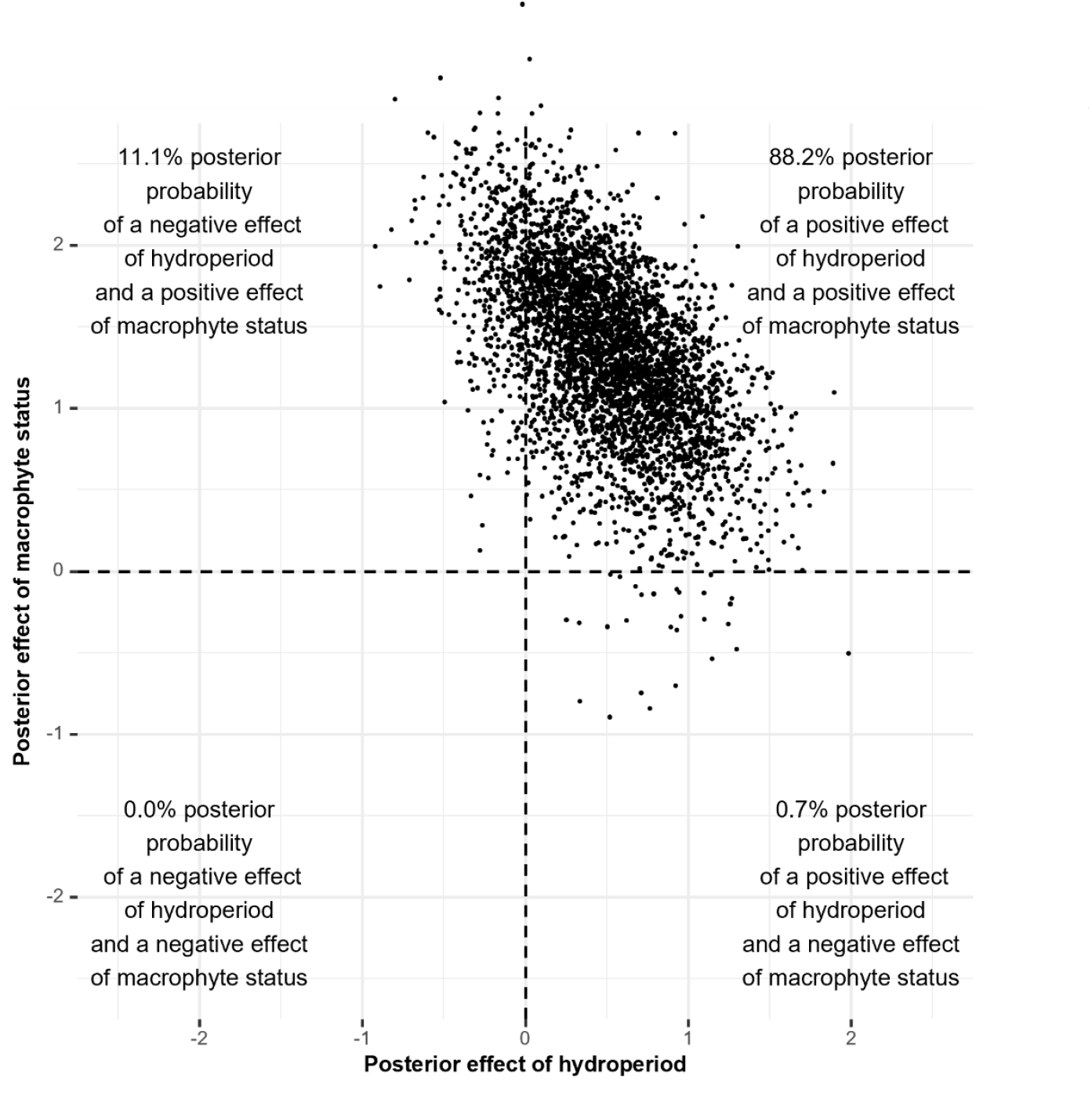
Joint posterior distribution for the effect of hydroperiod and macrophyte status on the species accumulation rates of the studied ponds. Points represent individual posterior draws. Vertical and horizontal lines represent the lines of no-effect, and categorize the two-dimensional space into four quadrants, each implying different biological conclusions, as clarified through corner texts.

In terms of species composition, a clear nested pattern was observed during the last year of the study period, with species-poor ponds generally hosting a subset of the species present in species-rich ponds. Nestedness was highly significant under all considered null model schemes (observed NODF = 68.03; p < 0.001), except for the most conservative one (p = 0.792). In total, 16 cladoceran species colonized the newly created ponds during the study period. *Daphnia obtusa* (Kurz, 1874) and *Chydorus sphaericus* (O.F. Müller, 1776) were by far the most widespread species, which colonized the vast majority of the ponds within the first three years of the pondscapes’ existence. *D. obtusa* emerged as the most successful colonizer, colonizing most ponds within the first six months after construction (Fig. 6). Nevertheless, colonization of *D. obtusa* in BOF7 was delayed until the third year, coinciding with a shift in local conditions, as this pond started retaining water for longer periods from that moment onwards (Janssens, pers. obs.). Overall, however, while being a successful colonizer, *D. obtusa* became less widespread as ponds age, i.e. during the third year of the study period (Supplementary information Table S1.2). Other fast-dispersing species included *Bosmina longirostris* (O.F. Müller, 1785) and *Daphnia ambigua* (Scourfield, 1947), although their occurrences were transient and typically observed only during a single sampling event.

**Fig. 6:**
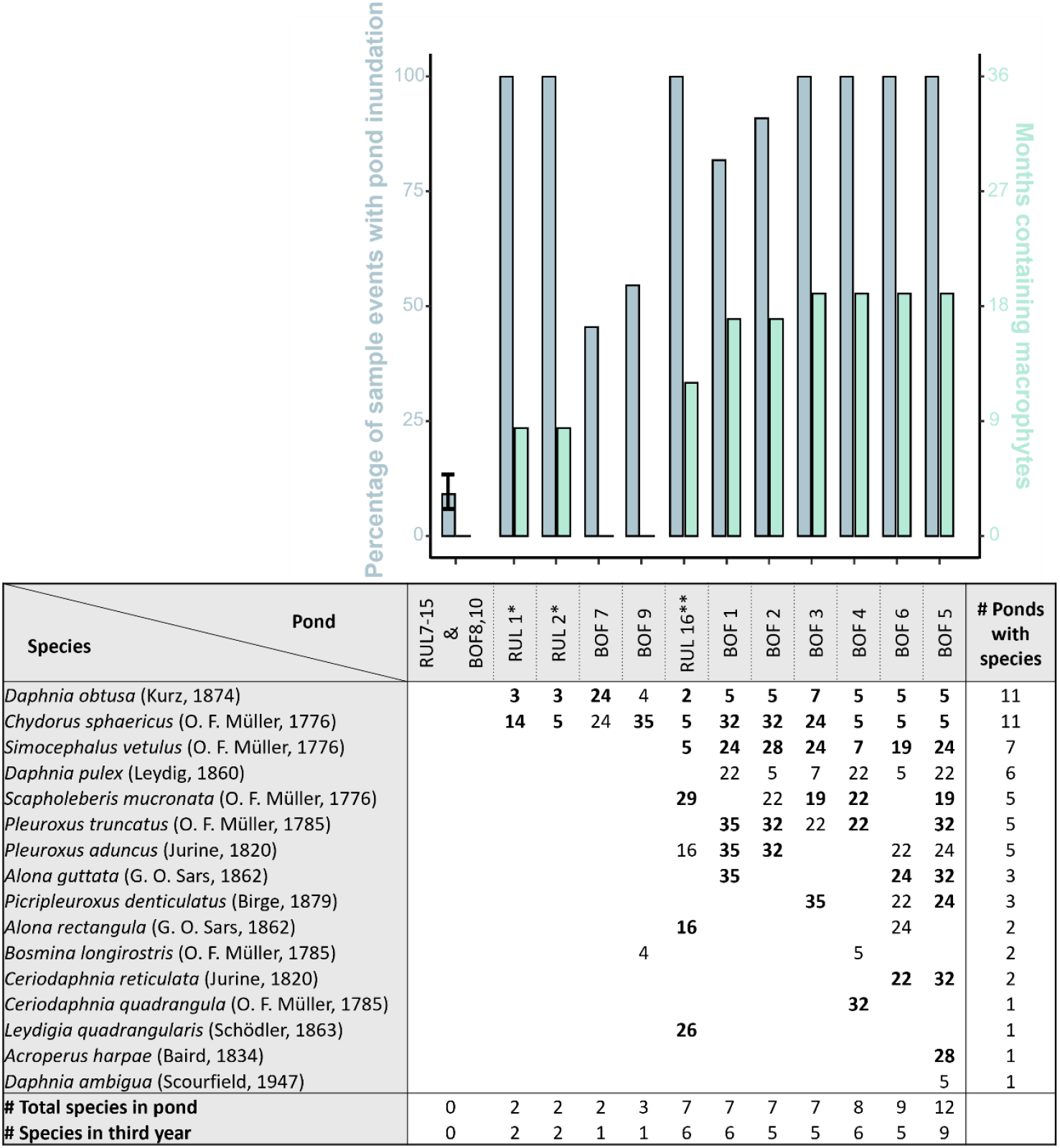
Upper: Barplot depicting the percentage of sampling moment that a given pond was inundated (blue) and the amount of months since creation that a pond contained macrophytes (green). Lower: Month of first species detection in each newly created pond (ponds are ranked based upon their cumulative species richness). Species still present in the third year of a pond’s existence are shown in bold. (*RUL1 and RUL2 were constructed in May 2023; **RUL16 in June 2022, these ponds were followed up until the pondscape was three years old, until they were 19 and 30 months old respectively)

In contrast to *D. obtusa, C. sphaericus* exhibited a more gradual colonization pattern, with most ponds only being colonized during their third year of existence. Despite its eventual widespread occurrence in the set of investigated ponds, its initial establishment is considerably slower compared to that of other cladoceran species. Also other chydorid species tended to colonize the ponds only in the last year. The increase in species richness observed during the second and third year was indeed primarily driven by the arrival of chydorid species (genera: *Acroperus, Alona, Leydigia, Pleuroxus, Picripleuroxus*), in addition to *Simocephalus vetulus* (O.F. Müller, 1776) and *Ceriodaphnia* species. This increase in species richness was most pronounced in ponds that were more permanent and contained macrophytes (Fig. 5).

## Discussion

Our results confirm earlier studies that some species, notably *D. obtusa*, are fast in colonizing ponds and that chydorids are slower in colonizing ponds, even though some species such as *Chydorus sphaericus* eventually become widespread (Louette & De Meester 2005). We also show that species accumulation trajectories differ among ponds and that the rate of species accumulation is higher in permanent ponds in which macrophyte vegetations establish.

The two most widespread species at the end of the study period (i.e. after three years) are *D. obtusa* and *C. sphaericus*. Both species were observed in fifteen of the studied ponds. But while the first observation of *D. obtusa* was in all but two ponds within the first seven months of the ponds’ existence, *C. sphaericus* was only observed in the second year or later in ten out of fifteen ponds. This observation is in line with Louette and De Meester (2005), who also observed that *D. obtusa* was the dominant first colonizer in a different set of ponds in a different region, whereas chydorids colonized those ponds later in time. This is striking, given that *D. obtusa* is not a particularly common species in general surveys (Louette et al., 2007). This suggests a high dispersal ability, but it may also reflect a capacity of *D. obtusa* to deal with unstable conditions in newly created ponds. Our data indeed show that the environmental characteristics of newly created ponds are highly variable in the first year of their existence, and that conditions converge in the second and third year. If *D. obtusa* is better able than most other species to survive in very young ponds, then our observations might reflect establishment success rather than colonization dynamics. *D. obtusa* has a short generation time and reproduces quickly compared to many other *Daphnia* species (Milbrink et al., 2003), and this might contribute to his success as early colonizer. Notably, *D. obtusa* becomes less widespread in the third year of our study. As ponds age, environmental pond conditions stabilize and more species colonize the ponds, and *D. obtusa* seems to be less successful in maintaining detectable populations. This again was also observed by Louette and De Meester (2005) and Louette et al. (2008). This consistency across studies in different regions suggests a general pattern, in which *D. obtusa* might behave as a so-called fugitive species that is very successful in early colonization and rapid population growth in young habitats and can occur in the landscape thanks to this effective dispersal even though it is weak in competition (Tilman et al., 1994).

There are a number of chydorid species, *Simocephalus vetulus* and *Scapholebreris. mucronata* (O.F. Müller, 1776), that occurred in multiple ponds, but were only first observed in the second or third year of a pond’s existence. This second wave of colonization coincided with the establishment of macrophytes in the ponds, which happened in month 18 or later. These species tend indeed to be either macrophyte-associated (chydorids, *Simocephalus*) or prefer sheltered waters, typically provided by emergent or floating-leaved vegetation (the hyponeustic species *Scapholeberis mucronata*). This suggests that the association that we observed between the development of macrophytes and the rate of colonization of the ponds by zooplankton reflects a causal relationship, where the presence of macrophytes increases cladoceran richness by providing greater habitat complexity and niche availability (Choi et al., 2014; Declerck et al., 2007). In a recent study by Peso et al. (2025), contrasting young and old ponds in survey data, it was observed that older ponds have higher species richness than younger ponds, but that the difference is especially striking in older ponds with a well-established vegetation of macrophytes.

Macrophytes were an important factor explaining variation in the increase of species richness between pondscapes and between ponds. Macrophytes facilitate the increase in cladoceran species richness through the establishment of multiple vegetation-associated taxa. A considerable proportion of the species present in the third year were indeed macrophyte-associated taxa, such as chydorids (*Acroperus, Alona, Leydigia, Pleuroxus, Picripleuroxus*), *S. vetulus*, and *Ceriodaphnia* spp. (Bledzki & Rybak, 2016). Given the strong correlation between macrophyte establishment and hydroperiod, however, hydroperiod should also be considered when interpreting patterns of species accumulation. Earlier studies have documented a central role of hydroperiod in structuring cladoceran communities (Coccia et al., 2024). Variation in hydroperiod of newly created ponds is therefore likely to, over-time, promote community divergence. This underscores the value of incorporating ponds with differing hydroperiod regimes within a pondscape in the context of nature restoration, as it contributes to an increase in overall cladoceran diversity at the regional level.

In our study macrophytes and hydroperiod are strongly colinear, which does not enable us to reliably assess their unique effects. Given that the later wave of colonization involved cladoceran species for which their association with macrophytes is very well documented, it seems logic to interpret our data as indicative of the importance of macrophytes, in line with the importance of well-developed macrophyte vegetation in determining the diversity of young ponds in earlier work (Peso et al., 2025). It is therefore difficult to derive strong guidelines on the importance of hydroperiod variation in nature restoration projects. Yet, given the unique species associated with temporary ponds (Céréghino et al., 2007; Williams, 1997), it is likely that the creation of variation in hydroperiod in the design of pondscapes will increase regional diversity of newly established pondscapes. In doing so, it will be crucial to consider climate warming in designing new pondscapes, given its important impact on pond hydroperiod (Brooks, 2009; Reid et al., 2019)

Spatial isolation did not have a statistically supported effect on species accumulation in our study, yet the estimated slopes suggest a weak tendency for ponds located closer to existing regional ponds to accumulate species at a slightly higher rate. A potential influence of distance aligns with expectations that dispersal from nearby source habitats can facilitate early community assembly, and earlier work indicated that the regional species richness affects colonization dynamics (Louette & De Meester 2005). Given the limited number of ponds included in the present study and the lack of variation in the distance to the closest regional pond, our ability to detect an effect of distance was low,.

## Conclusion

Our results show that *D. obtusa* is a fast colonizing cladoceran species that successfully establishes in newly created ponds already in first few months after pond creation. After three years, its dominance declines when additional species arrive. This pattern suggests that *D. obtusa* acts as a fugitive species that thrives in unstable, early-succession pond conditions where it temporarily can be the dominant species, before other species such as *C. sphaericus* and *S. vetulus*. Our study also confirms that macrophyte presence is a key factor explaining variation in species richness build-up between pondscapes and among individual ponds, supporting the additional colonization by macrophyte-associated zooplankton taxa. The findings of our study highlight the role of macrophyte presence in shaping cladoceran species richness build-up during early pond succession, which offers valuable insights for designing effective pond creation supporting rapid cladoceran species accumulation.

## Acknowledgements

We thank Foundation Elzéard and its staff for their cooperation and for granting us permission to study the ponds they have created. We specifically thank Robert Delbroek for his assistance throughout the study period. We thank everyone participating in fieldwork, and in particular Louis-Marie Le Fer and Ditte Rens for their contribution. This work was financially supported by Research Foundation Flanders (FWO, grant number: G0A3M24N), the PONDERFUL project that is funded by the European Union’s Horizon 2020 research and innovation program under grant agreement No. 869296, Biodiversa+ project TRANSPONDER, and KU Leuven Research Council project C16/2023/03. R.W. acknowledges a KU Leuven PhD scholarship (grant number DB/22/007/bm).

## Data availability statement

All data and code for these analyses are available on Github: https://github.com/BramJanssens1/Cladoceran_richness_accumulation

