## Supplementary Information for "Cladoceran diversity build-up in newly created pondscapes"

°Shared first authorship

**ORCID**

Bram Janssens: 0009-0005-5734-1716

Robby Wijns: 0000-0002-9548-4030

Maxime Fajgenblat: 0000-0002-2233-1527

Pieter Lemmens: 0000-0002-3135-9724

Thomas Neyens: 0000-0003-2364-7555

Luc De Meester: 0000-0001-5433-6843

**Supplementary information**

**SI1 Supplementary information main analyses**

***Table S1.1:*** *Ponds with their respective distance(m) to the closest regional pond (i.e. ponds that existed prior to the habitat creation project, that might have served as source habitats for the colonization of newly created ponds), maximal depth (cm), and maximal surface area (m^2^) during the sampling period.*

| **PondID** | **Distance to nearest regional pond (m)** | **Maximum depth (cm)** | **Maximum pond surface area (m^2^)** |
| --- | --- | --- | --- |
| RUL1 | 951.05 | 180 | 27.0 |
| RUL2 | 949.56 | 100 | 25.6 |
| RUL16 | 721.00 | 43 | 3.2 |
| BOF1 | 317.36 | 32 | 3.0 |
| BOF2 | 322.23 | 48 | 2.4 |
| BOF3 | 301.88 | 118 | 4.2 |
| BOF4 | 148.05 | 96 | 16.5 |
| BOF5 | 146.91 | 99 | 18.0 |
| BOF6 | 152.43 | 104 | 17.4 |
| BOF7 | 89.60 | 68 | 13.5 |
| BOF9 | 230.78 | 45 | 2.5 |

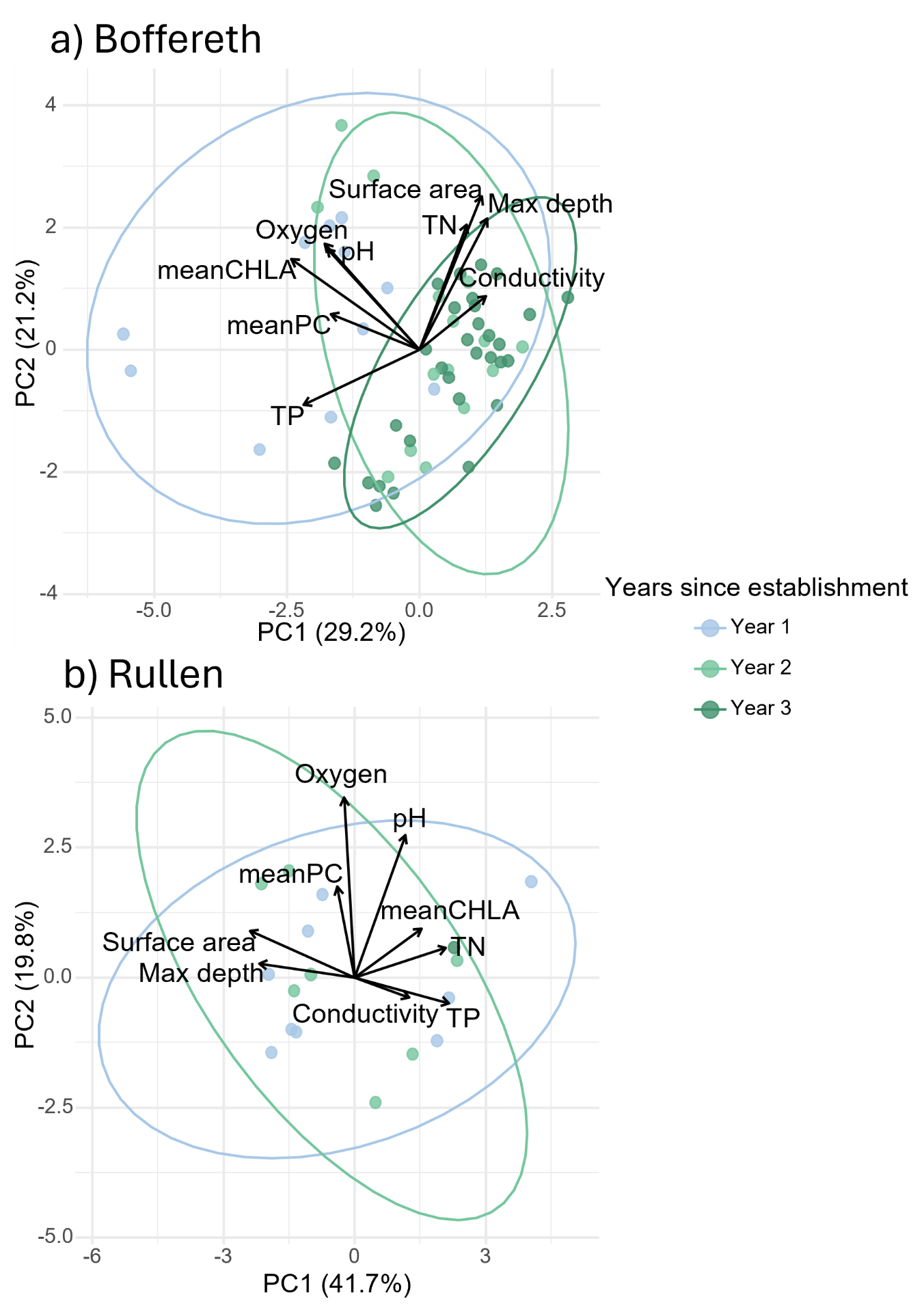

***Fig. S1.1****: PCA ordination plot based on the local environmental and morphometric characteristics of the newly created ponds in the Boffereth (a) and the Rullen (b) region separately, during the first three years of their existence. Different years are visualized using ellipses reflecting 0.95 confidence intervals.*

***Table S1.2:*** *Presence (1) and absence (0) of* Daphnia obtusa *across 11 sampling moments during the first three years of the pondscapes' existence. Ponds that had not yet been created during a particular sampling moment are indicated with an "x", while ponds that were present but dry are represented by a grey square. The last row reflects the percentage of water-containing ponds in which* D. obtusa *was detected at each time point.*
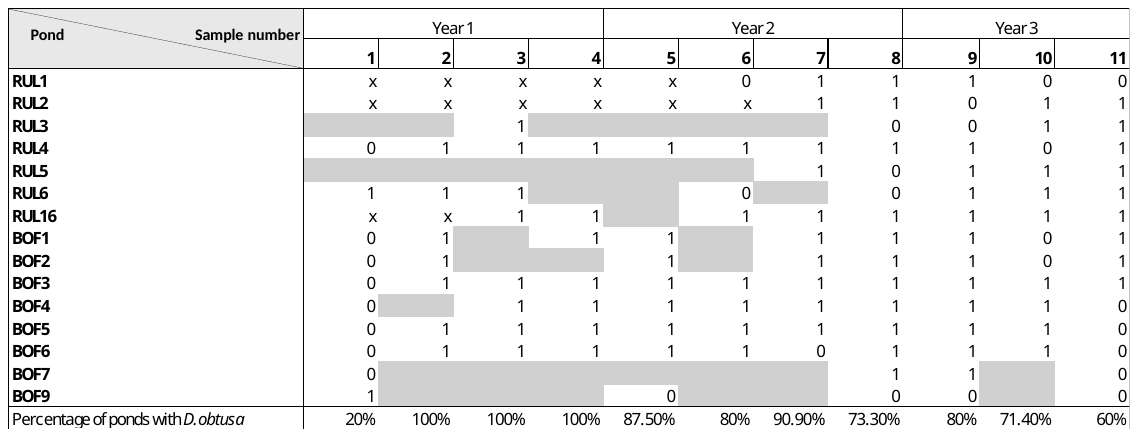

**SI2: Accidentally inoculated ponds**

During construction four ponds of the Rullen pondscapes were filled up using water from a well that we later sampled and contained *Daphnia obtusa, Chydorus sphaericus* and *Simocephalus vetulus*. Due to the potential effects of this artificial inoculation on long term community assembly, were these ponds not included in the main analysis of the manuscript. Identical analyses including the inoculated ponds are visible below (Fig. S2.1-S2.5).

Despite the observed difference in accumulation rate between ponds within pondscapes (Fig. S2.1), we did not find strong statistical support for a direct effect of pondscape identity per se (76.5% posterior probability of an effect). An average pond (i.e. conditional on average characteristics, except for pondscape identity) in Rullen and Boffereth displays an estimated accumulation rate of 2.1 species per year (95% CrI [1.4; 2.7]) and 1.7 species per year (95% CrI [0.9; 2.5]), respectively (Fig. S2.3). We found strong statistical support that well-vegetated ponds feature higher accumulation rates than less-vegetated ponds (99.7% posterior probability of an effect). The average well-vegetated pond (highest recorded coverage of emersed or submerged macrophytes >25%) displays an estimated accumulation rate of 2.5 species per year (95% CrI [1.3; 3.8]), while the average less-vegetated pond displays an estimated accumulation rate of 1.3 species per year (95% CrI [0.0; 2.5]) (Fig. S2.3). This corresponds to a difference in accumulation rate of 1.3 species per year (95% CrI [0.5; 2.1]). We also found strong statistical support for a positive effect of hydroperiod (98.5% posterior probability). The average pond containing water during all sampling moments displays an estimated accumulation rate of 2.1 species per year (95% CrI [0.8; 3.4]), while the average pond containing water during half of the sampling moments displays an estimated accumulation rate of 1.4 species per year (95% CrI [0.2; 2.5]) (Fig. S2.3). This corresponds to an estimated difference in accumulation rate of 0.7 species per year (95% CrI [0.1; 1.4]). We found the joint posterior distribution of macrophyte status and hydroperiod to be negatively associated, reflecting that macrophyte vegetations are better developed in ponds with longer than with shorter hydroperiods (Fig. S2.4). As a result of this collinearity, the model cannot fully disentangle the direct effect of macrophyte status and hydroperiod at a 95% certainty threshold. While we found strong and moderately strong statistical support for the marginal effect of macrophytes and hydroperiod, respectively, the available data does not allow to fully isolate the effect of macrophyte status, leading to a more uncertain estimate and moderately strong statistical support. We found 98.2% posterior probability that both variables are directly related to the species accumulation rate, but only a 0.3% posterior probability for a pure effect of macrophyte  (i.e., excluding the shared effect with hydroperiod), and a 1.5% posterior probability for a pure effect of hydroperiod (i.e., excluding the shared effect with macrophyte status; Fig. S2.4). We did not find any evidence for the influence of local environmental conditions as summarized by the temporally averaged first and second principal component coordinates (70.6% and 89.8% posterior probability of an effect, respectively) (Fig. S2.3). We did find strong statistical support for the effect of distance to the closest existing, regional pond (92.6% posterior probability of an effect).

Interestingly, these four ponds (RUL3–6) did not show any difference in species richness build-up compared to the other ponds of the Rullen site. The median time until first colonization in non-contaminated ponds did not differ from that of the contaminated ponds for *D. obtusa* (5 months vs. 6 months) and was longer for *C. sphaericus* (19 months vs. 32 months). While *S. vetulus* was only observed in one of the contaminated ponds, it was more prevalent in non-contaminated ponds (Fig. S2.5).

The dynamics in these four ponds that were accidentally contaminated illustrate the importance of environmental sorting in the observed colonization dynamics. Despite their likely introduction when the ponds were artificially filled with well water, *C. sphaericus* and *S. vetulus* failed to establish and did not alter the ponds future cladoceran build-up compared to non-contaminated ponds. This reflects that there are important pressures that the local environment is exerting on these species, unabling to establish after inoculation. The observation that *D. obtusa* is capable to establish in these systems emphasizes it being adapted to establish in highly unstable systems.

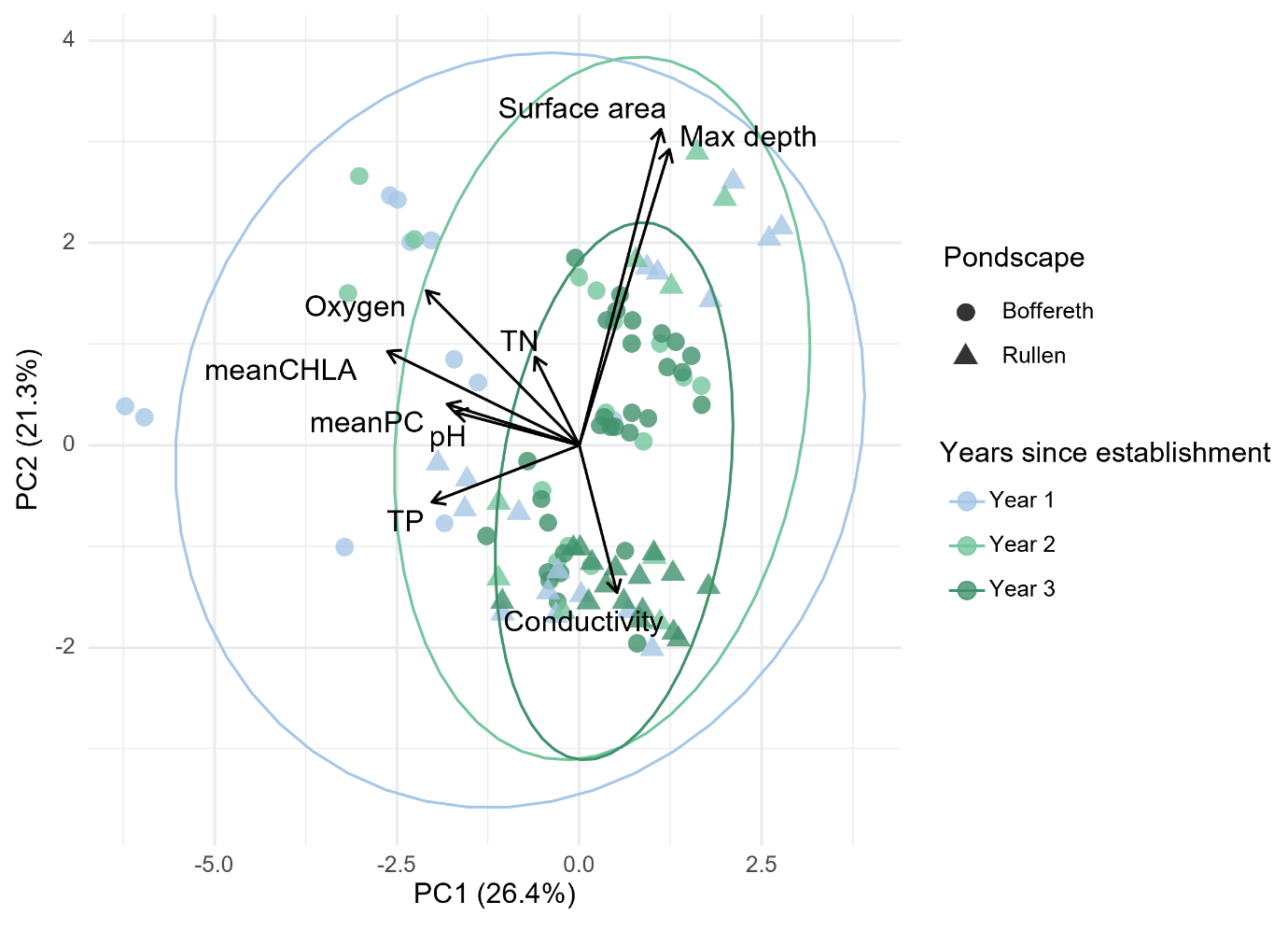

***Fig. S2.1:*** *PCA ordination plot based on the environmental pond characteristics for each sampling moment, maximum depth and surface area during the first three years of their existence. Different years are highlighted using different 95% confidence ellipses. Pondscapes are visualized by different symbols (Boffereth: circles; Rullen: triangles) (including the accidental inoculated ponds).*

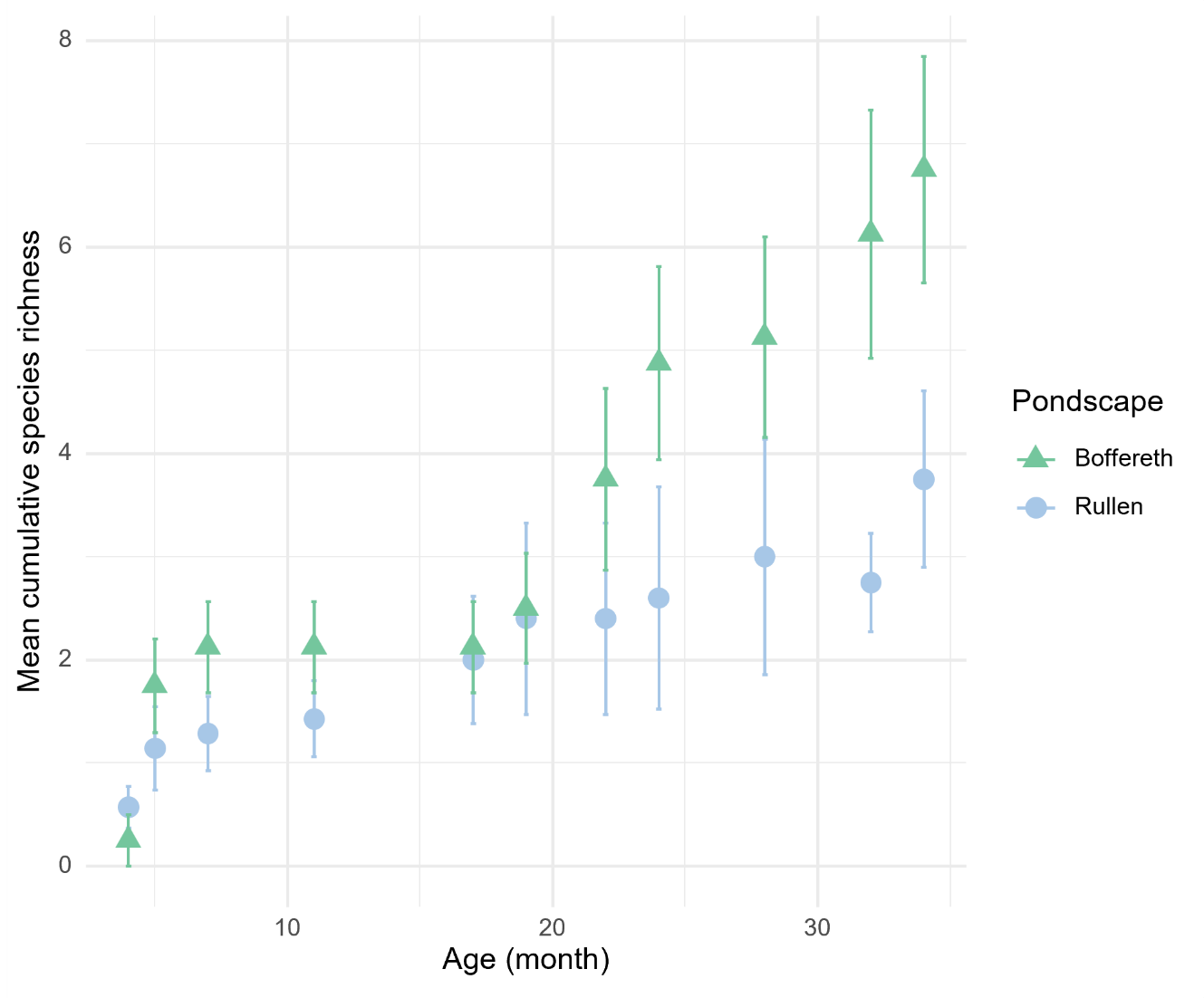

***Fig. S2.2:*** *Average cumulative species richness of cladocerans during the first three years after pond creation for each newly created pondscapes: Boffereth (green triangles) and Rullen (blue circles) (including the accidental inoculated ponds). Values are averaged across ponds per pondscapes. Error bars indicate the 95% standard error.*

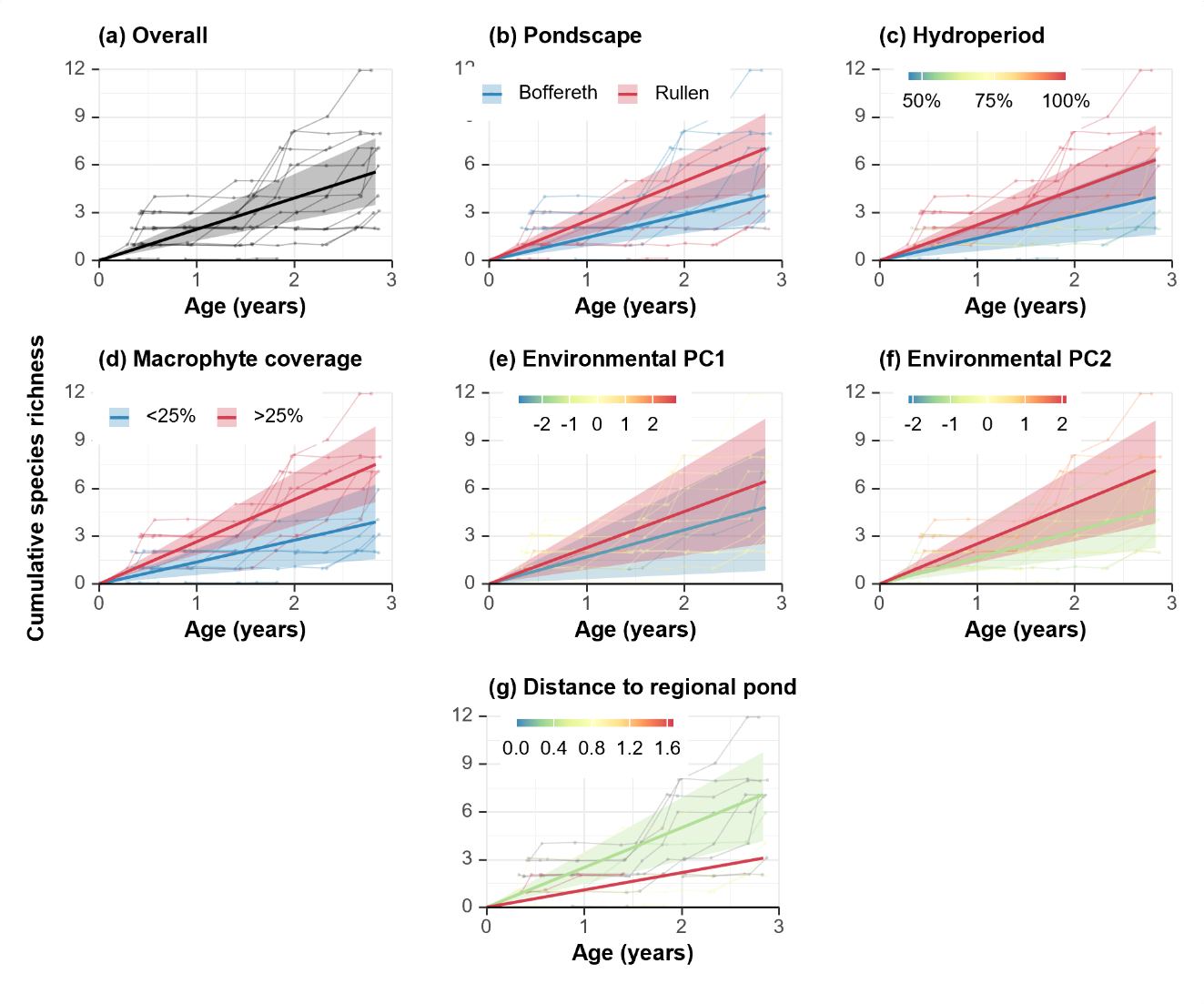

***Fig. S2.3:*** *Estimated, conditional linear species accumulation trajectories as a function of pond age, all other conditions being held at the average value across the dataset. Individual points represent observed cumulative species richnes data across ponds (including the accidental inoculated ponds) and time points, with points belonging to the same pond being joined by thin lines. Estimated linear species accumulation trajectories are shown using thick lines, with shaded areas representing 95% credible intervals. (a) Estimated trajectory for the average pond. (b) Estimated trajectory for the average pond in the pondscapes Boffereth and Rullen. (c) Estimated trajectory for the average permanent and temporary pond. (d) Estimated trajectory for the average well- and less-vegetated pond. (e) Estimated trajectory for the average pond with a low and high local environmental condition as represented by the PC1 coordinate. (f) Estimated trajectory for the average pond with a low and high local environmental condition as represented by the PC2 coordinate. (g) Estimated trajectory for the average pond with a low and high distance to the closest regional pond.*

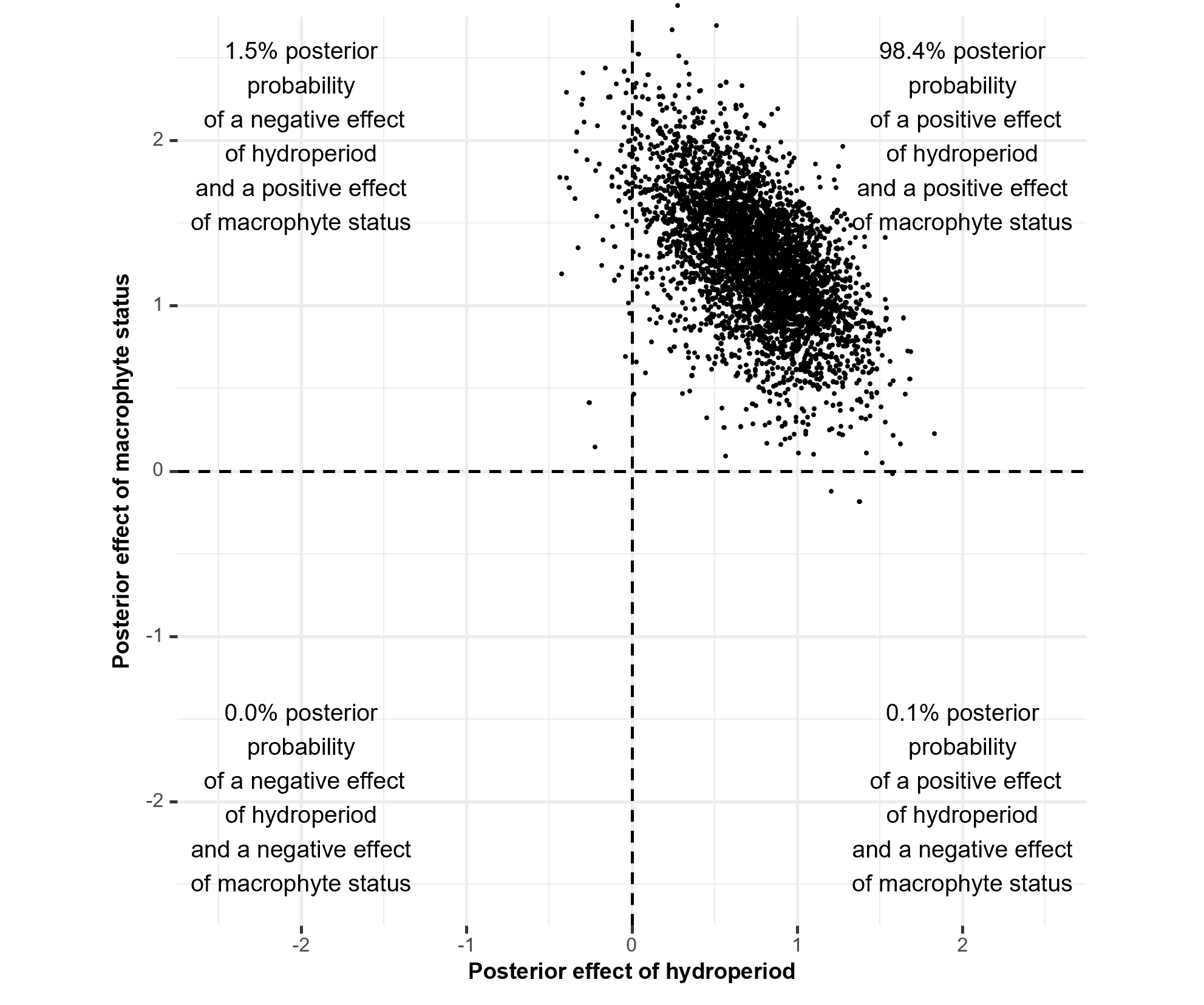

***Fig. S2.4:*** *Joint posterior distribution for the effect of hydroperiod and macrophyte status on the species accumulation rates of the studied ponds (including the accidental inoculated ponds). Points represent individual posterior draws. Vertical and horizontal lines represent the lines of no-effect, and categorize the two-dimensional space into four quadrants, each implying different biological conclusions, as clarified through corner texts.*

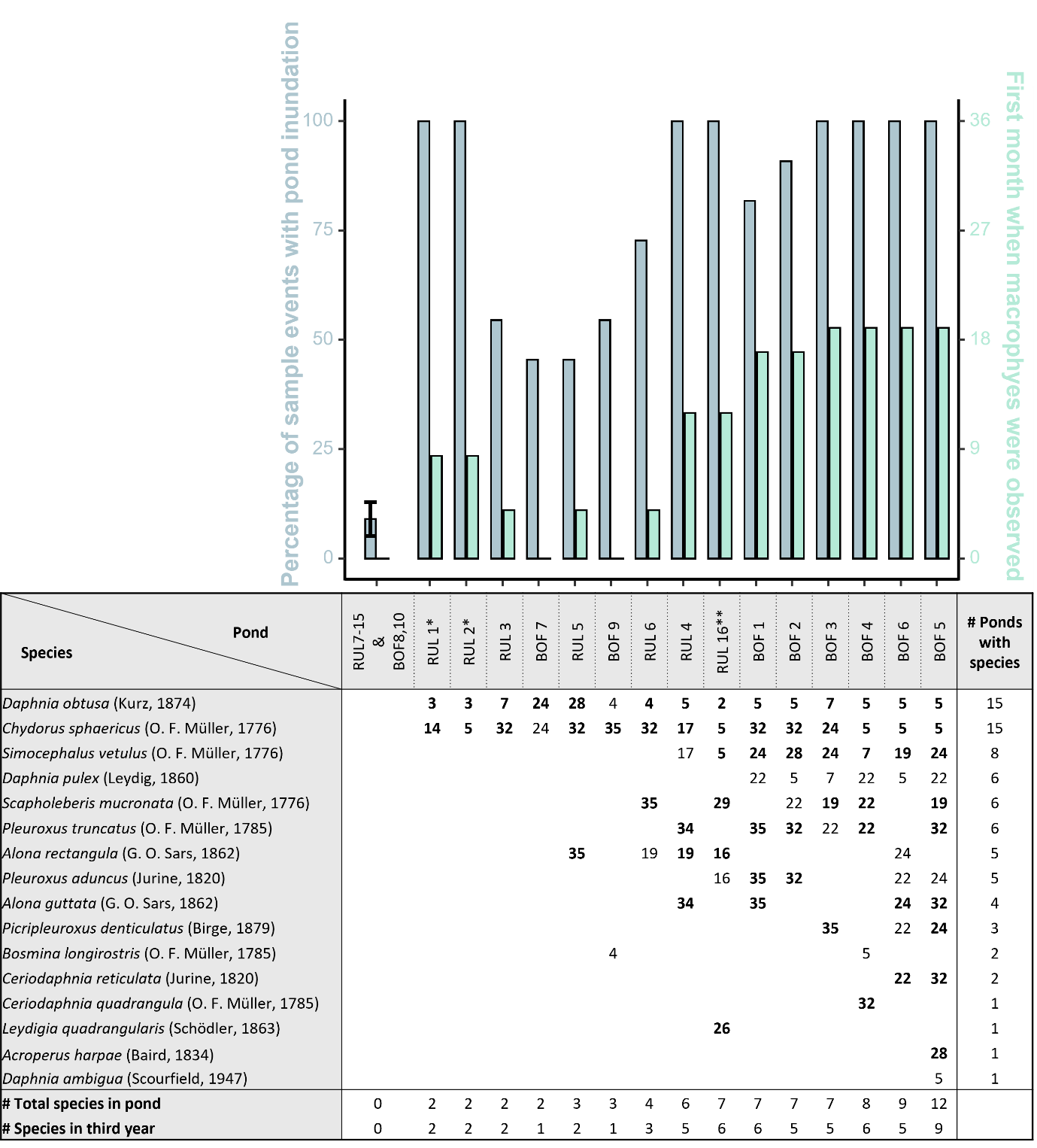

***Fig. S2.5:*** *Upper: Barplot depicting the percentage of sampling moment that a given pond was inundated (blue) and the amount of months since creation that a pond contained macrophytes (green) (including the accidental inoculated ponds). Lower: Month of first species detection in each newly created pond (ponds are ranked based upon their cumulative species richness). Species still present in the third year of a pond’s existence are shown in bold. (*RUL1 and RUL2 were constructed in May 2023; **RUL16 in June 2022, these ponds were followed up until the pondscape was three years old, until they were 19 and 30 months old respectively)*
